# Mitochondrially targeted proximity biotinylation and proteomic analysis in *Plasmodium falciparum*

**DOI:** 10.1101/2022.05.30.494025

**Authors:** Ian M. Lamb, Kelly T. Rios, Anurag Shukla, Avantika I. Ahiya, Joanne Morrisey, Joshua C. Mell, Scott E. Lindner, Michael W. Mather, Akhil B. Vaidya

## Abstract

Despite ongoing efforts to control malaria infection, progress in lowering the number of deaths and infections appears to have stalled. The continued high incidence of malaria infection and mortality is in part due to emergence of parasites resistant to frontline antimalarials. This highlights the need for continued identification of novel protein drug targets. Mitochondrial functions in *Plasmodium falciparum*, the deadliest species of human malaria parasite, are targets of validated antimalarials including atovaquone and proguanil (Malarone). Thus, there has been great interest in identifying other essential mitochondrial proteins as candidates for novel drug targets. Garnering an increased understanding of the proteomic landscape inside the *P. falciparum* mitochondrion will also allow us to learn about the basic biology housed within this unique organelle. We employed a proximity biotinylation technique and mass spectrometry to identify novel *P. falciparum* proteins putatively targeted to the mitochondrion. We fused the leader sequence of a mitochondrially targeted chaperone, Hsp60, to the promiscuous biotin ligase TurboID. Through these experiments, we generated a list of 122 “putative mitochondrial” proteins. To verify whether these proteins were indeed mitochondrial, we chose five candidate proteins of interest for localization studies using ectopic expression and tagging of each full-length protein. This allowed us to localize four candidate proteins of unknown function to the mitochondrion, three of which have previously been assessed to be essential. We suggest that phenotypic characterization of these and other proteins from this list of 122 could be fruitful in understanding the basic mitochondrial biology of these parasites and aid antimalarial drug discovery efforts.

## Introduction

The complex life cycle of *P. falciparum* takes place in two host species: humans and mosquitoes. The asexual blood stage causes malaria symptoms in humans and the sexual stage in mosquitoes is essential for malaria transmission. Asexual parasites rely on glycolysis for energy generation with minimal contributions from mitochondrial oxidative phosphorylation. Furthermore, the canonical tricarboxylic acid (TCA) cycle is not essential at this stage but is essential during the insect stages of the parasites [1, 2]. Although not significantly active in energy generation via respiration in asexual blood stage parasites, this “minimal” single mitochondrion serves other essential functions, such as supporting pyrimidine biosynthesis, and is a validated target of antimalarials that target the asexual stage [3–6]. The 6 kb mitochondrial DNA (mtDNA) of malaria parasites encodes only three proteins (subunits 1 and 3 of cytochrome *c* oxidase and cytochrome *b*), which are essential proteins of the mitochondrial electron transport chain (mtETC), and ribosomal RNAs [7–10]. However, >400 nuclear-encoded proteins are believed to be imported into the mitochondrion to perform other essential functions such as iron sulfur cluster synthesis, as well as maintenance, transcription, and translation of mtDNA. Because of the therapeutic success of targeting the mitochondrial electron transport chain (mtETC) in asexual blood stage parasites, such as inhibition of the parasite’s cytochrome *bc_1_* complex by the antimalarial atovaquone, there has been great interest to identify other essential mitochondrial proteins [11, 12]. Toward this effort, a complete and experimentally driven mitochondrial proteome of *P. falciparum* could provide guidance for drug discovery efforts.

A common proximity biotinylation technique uses a promiscuous biotin ligase fused to a bait protein to induce biotinylation of nearby proteins, followed by pull down and liquid chromatography coupled to tandem mass spectrometry (LC-MS/MS) to infer their identify from peptide sequences [13]. Recently, a highly efficient biotin ligase “TurboID” was constructed to allow shorter labeling periods and reduced nonspecific labeling compared to the original BirA* variant used in BioID [14]. Another approach to improve the specificity of labeling would be to express the ligase in a conditional manner. We have taken advantage of conditional TetR-DOZI expression system that is controlled by anhydrotetracycline (aTc) [15], in which a protein is translated in the presence of this small molecule, to investigate the mitochondrial proteome in *P. falciparum*. We generated a parasite line for controlled labeling using a construct in which the leader sequence of the mitochondrially targeted chaperone Hsp60 [16] was fused to TurboID and expressed conditionally in the presence of aTc. The goal of this approach was to minimize labeling of cytosolic proteins by the biotin ligase on its way to be imported into the mitochondrion by shutting down its expression through the removal of aTc, and then carry out labeling by the ligase already imported into the mitochondrion prior to and immediately after the removal of aTc. Here, we describe unexpected limitations of this approach that may be due to unique aspects of mitochondrial protein import in *P. falciparum*. We also describe our approaches to work around these challenges that led to identification of 122 putative mitochondrial proteins and go on to validate mitochondrial targeting of some of these targets by generating transgenic parasites expressing tagged versions of these proteins.

## Results

### Generation of a parasite line conditionally expressing Hsp60 leader-TurboID-3xHA

We constructed a plasmid in which the 69 amino acid N-terminal leader sequence of mitochondrially-localized chaperone Hsp60 (Hsp60L) was fused to the promiscuous biotin ligase TurboID with three C-terminal copies of hemagglutinin epitope (3xHA) tag (Fig 1A). To integrate the plasmid at a nonessential genomic site, we employed a mycobacteriophage *Bxb1* integrase system adapted for use in *P. falciparum* [17]. In this system, a chromosomal *attB* locus is introduced into the non-essential *GLP3* gene. The *attP* site on the incoming plasmid then recombines with the *attB* locus, allowing stable expression of the gene-of-interest from the genomic site (Fig 1A). Correct integration was confirmed by diagnostic PCR with primer annealing sites depicted (Fig 1B). The trans-gene contained 8 copies of TetR binding aptamers within its 3’ untranslated region (UTR) that are recognized by the TetR-DOZI fusion protein, which is also encoded by this plasmid. In the absence of aTc, TetR-DOZI binds to the aptamers in the UTR of the target mRNA and represses its translation, whereas the addition of aTc displaces TetR-DOZI from the aptamers releasing the mRNA to be translated [15]. The rationale for this approach was to reduce biotinylation of cytosolic proteins by the Hsp60L-TurboID protein in transit to the mitochondrion by turning off translation of the trans-gene by removal of aTc (Fig 1C). We confirmed that the Hsp60L-Turbo-ID-3xHA fusion protein was significantly expressed only in the presence of aTc (Fig 1D).

**Fig 1.**
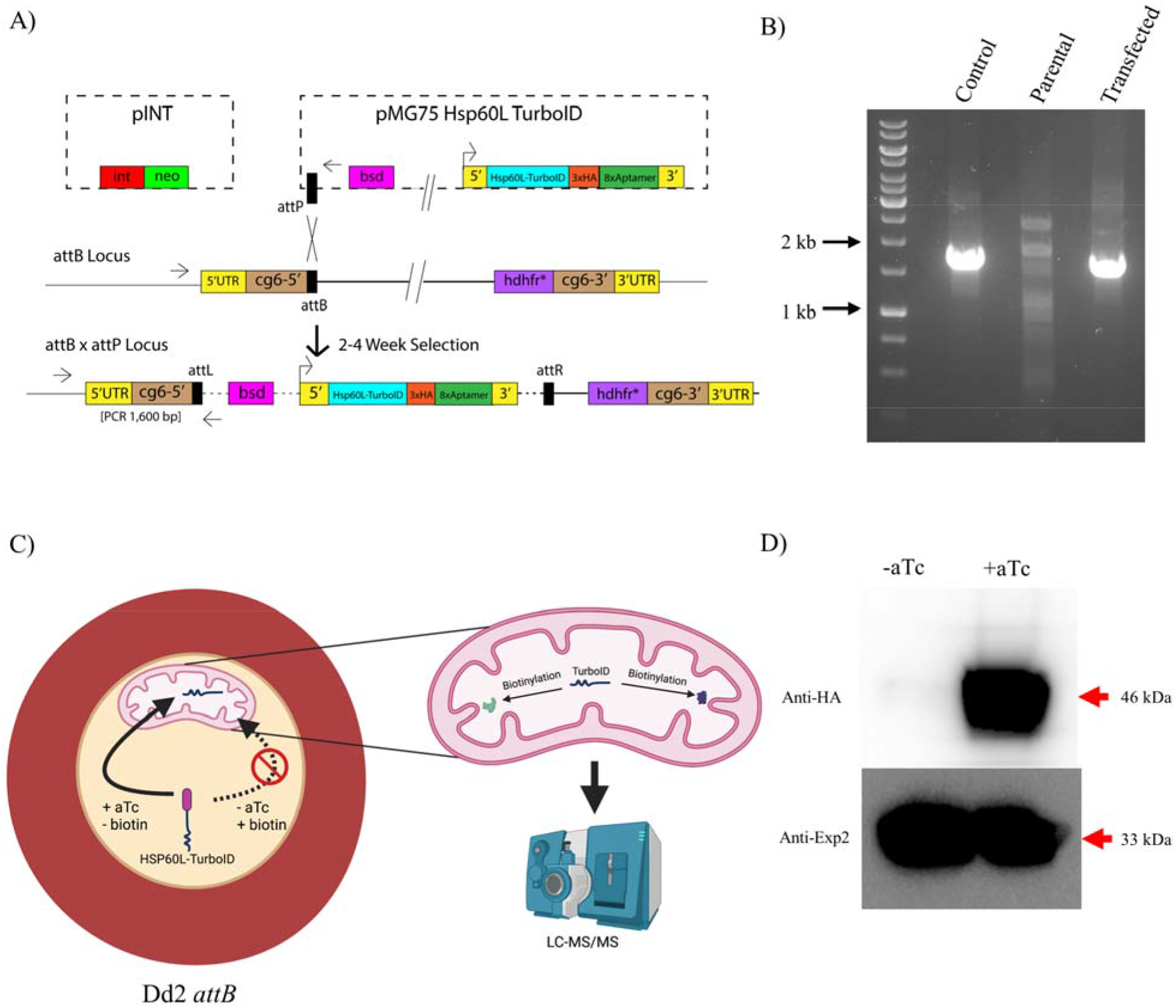
Generation of parasites expressing Hsp60L-TurboID regulated by the presence of aTc. A) Diagram of the plasmids used and mechanism of integration of Hsp60L-TurboID into Dd2 *attB* parasites. B) Agarose DNA gel image of PCR reactions confirming successful genomic integration of Hsp60L-TurboID. The PCR product obtained with DNA from the transgenic line runs at the expected size of ~1.6 kb. The DNA marker used is GeneRuler 1 kb DNA ladder (ThermoFisher Scientific). C) Schematic diagram of initial experimental design. Parasites grown in biotin free medium in the presence of aTc display robust targeting of Hsp60L-TurboID to the mitochondrial matrix. Upon aTc withdrawal, the fusion protein is knocked down and synthesized Hsp60L-TurboID proteins remaining in the cytosol are imported into the mitochondrion. Upon the addition of biotin, labelling of mitochondrial proteins accessible to the imported TurboID occurs. D) Western blot analysis of saponin-freed Hsp60L-TurboID parasites grown in +/- aTc conditions. PVDF membrane was probed with an anti-HA antibody with anti-Exp2 antibodies as a loading control.

To determine if TurboID was biotinylating parasite proteins *in vivo*, we incubated parasites in the presence or absence of aTc with 150 μM biotin for 2 or 4 hours (Fig 2A). We observed only minimal biotinylation of proteins under the aTc^-^ /biotin^+^ condition but robust biotinylation in the aTc^+^/biotin^+^ condition. Indirect immunofluorescence assays using an anti-HA antibody showed that Hsp60L-TurboID-3xHA was localized mainly to the parasite mitochondrion (Fig 2B). We also assessed whether the proteins biotinylated by TurboID were localized within the mitochondrion. To accomplish this, we incubated the parasites with biotin for 1 hour prior to fixation followed by detection with FITC-labeled streptavidin. As shown in Fig 2B, biotinylated proteins overlapped with MitoTracker staining. To determine if there is a fitness cost associated with non-physiological biotinylation of mitochondrial proteins, parasites were grown in complete, biotin-containing RPMI with or without aTc. As seen in Fig 2C growth of these transgenic parasites was indistinguishable whether or not aTc was included in the growth medium, showing no apparent fitness cost to the parasite.

**Fig 2.**
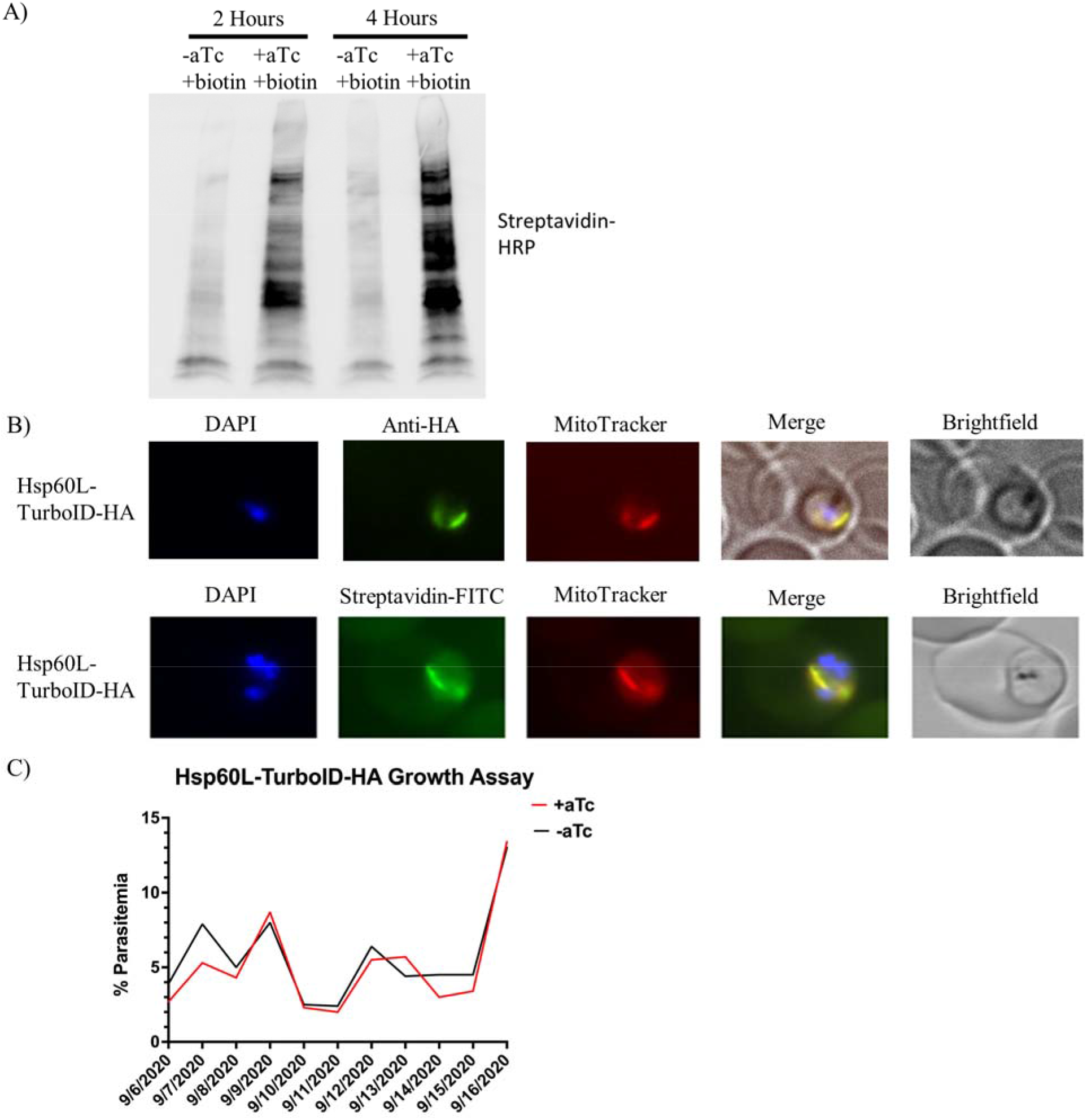
Characterization of pMG75BB-8XAptamer-TurboID-3xHA parasite line. A) Western blot analysis of equal numbers of parasites grown in aTc^-^/biotin^+^ or aTc^+^/biotin^+^ for a period of 2 or 4 hours. The blotted PVDF membrane was probed with streptavidin-HRP. Note the time-dependent increase in biotinylated proteins between hours 2 and 4 in the aTc^+^ biotin^+^ condition. B) Indirect immunofluorescence assay using an anti-HA antibody in aTc^+^ medium to confirm mitochondrial localization of Hsp60L-TurboID-3xHA and an immunofluorescence assay using a streptavidin-FITC conjugate. For this assay, parasites were grown in aTc^+^ medium and incubated with 150μM biotin for 1 hour. Note the overlap between biotinylated proteins and Mito-Tracker staining. C) Hsp60L-TurboID growth assay. Parasites were split equally to maintain healthy parasitemia. Giemsa-stained slides were made daily for 11 days. Parasitemia was determined each day by counting the number of infected red blood cells per 1,000 red blood cells per slide.

### Proximity biotinylation experiments targeting the P. falciparum mitochondrial matrix

Having established conditional expression of the transgene, we carried out a time course biotinylation experiment. Six flasks each containing 4×10^8^ trophozoite-stage parasites grown in biotin-free growth medium with aTc were used in the experiment. As shown in Fig 3A, aTc was washed out and the flasks were incubated in biotin-free medium for 3 h, at which point 150 mM biotin was added to 2 flasks, incubated for 1 h and harvested for protein isolation. These samples comprised replicate A and B of sample 1 (S1A and S1B). Five hours after removal of aTc, biotin was added to 2 other flasks and after 1 h of incubation, parasites were harvested (replicate A and B of Sample 2, S2A and S2B). Finally, at 7 h after the removal of aTc, biotin was added to the remaining 2 flasks and parasites were harvested following 1 h of incubation (Replicate A and B of Sample 3, S3A and S3B). A portion of the harvested proteins from each sample and replicate were used for SDS-PAGE and western blot analysis. As shown in Fig 3B, the HA-tagged protein product is present in two forms: a higher molecular weight band corresponding to full length Hsp60L-TurboID, and a lower one in which the Hsp60 mitochondrial targeting peptide was removed after import into the mitochondrion. We expected that, as the knockdown proceeded over time, we would observe a reduction of the higher molecular weight band and a relative increase in the lower molecular weight band. The rationale for this expectation is that once Hsp60L-TurboID was no longer being synthesized, proteins already synthesized would be imported into the mitochondrion over time. Surprisingly, the intensity of both bands diminished at the same rate (Fig 3B), suggesting either slow processing of the precursor by the mitochondrial processing peptidase after its import or retention of some the protein in the cytoplasm. To distinguish between these possibilities, we conducted LC-MS/MS analysis of each sample and its replicate. In the case of inefficient cleavage by the mitochondrial processing peptidase, we would expect to see reduced spectral counts of known cytosolic proteins as the knockdown proceeded over time, and relatively stable spectral of known mitochondrial proteins. However, we did not see any apparently significant differences in spectral counts across the entire time course of knockdown (Fig. 3C). Clearly, while the rationale for our approach of conditional expression of biotinylation enzyme appeared sound, the data suggest continued retention of a portion of this fusion protein in the cytoplasm.

**Fig 3.**
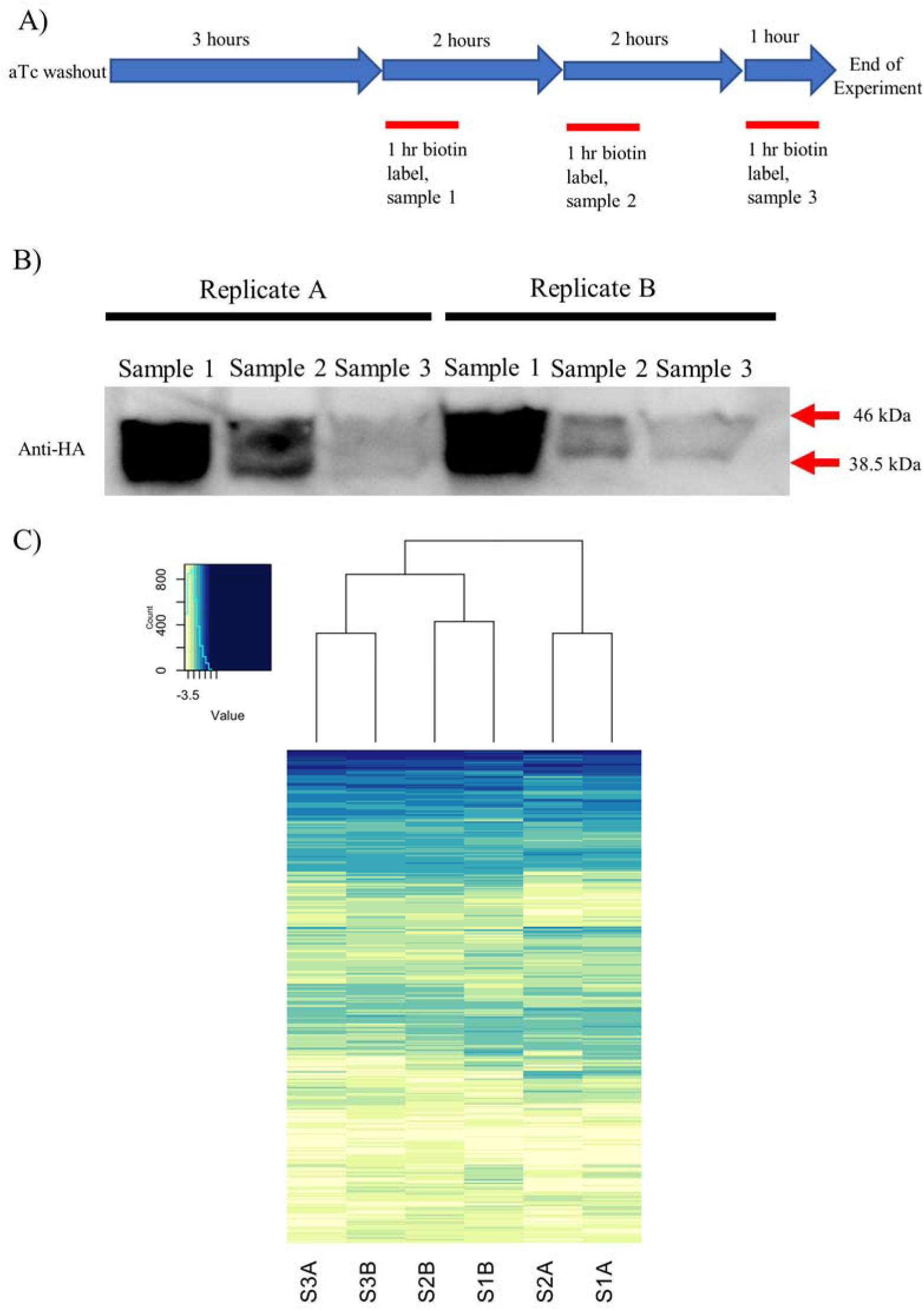
Protein biotinylation dynamics after different time periods of knockdown. A) Timeline of how these experiments were carried out (n = 2). B) Western blot analysis of Hsp60L-TurboID parasites knocked down for 4, 6, or 8 hours. Note that Hsp60L-TurboID-3xHA runs as a doublet. The higher molecular weight band corresponds to full-length Hsp60L-TurboID-3xHA. The lower molecular weight band probably corresponds to TurboID-3xHA that has been imported to the mitochondrion with concomitant removal of the Hsp60 targeting sequence. C) Heat map of the relative intensity of spectral counts for each protein identified by LC-MS/MS. Heat map represents 609 proteins with at least one spectral count in all samples. The scale is log_10_ (counts + pseudo-count), where pseudo-count = 1×10^e-4^ with the whitest color being 0. Drawing of row dendrograms and row labels is suppressed. Heat map was drawn using RStudio. Spectral count data derived from Trans Proteomic Pipeline analysis for each sample in this experiment is available in S1 Table.

Given this unexpected challenge, we abandoned the knockdown approach in favor of physical separation of “soluble” (cytoplasmic) proteins from “membrane” (i.e., organellar) proteins after biotin labeling using hypotonic lysis and differential centrifugation. The experimental setup is described in detail in the methods section. Western blot analysis of “soluble” and “membrane” fractions showed that while the soluble fraction contained only full-length Hsp60L-TurboID-3xHA, the membrane fraction contained both full-length and the processed species (Fig 4A), indicating higher mitochondrially targeted protein enrichment. We hypothesized that by comparing spectral counts of biotinylated proteins between these two fractions, we could identify the proteins biotinylated by imported TurboID-3xHA and thus more likely to be mitochondrial. We therefore conducted three biological replicates using this fractionation approach. In each experiment, the membrane and soluble fractions were generated and incubated with streptavidin-conjugated nanobeads, then washed extensively with a buffer containing 2% w/v SDS. Tryptic peptides from the bound proteins were analyzed by LC-MS/MS. As a control, isogenic parasites not incubated with biotin (“no biotin control”) were also generated using the same procedure. We analyzed proteins detected in each biological replicate of these experiments using the SAINT algorithm (Significance Analysis of INTeractome) to normalize spectral counts to protein length and the total spectra in each experiment, and to statistically identify high confidence hits in the membrane and soluble fractions [18] (Fig 4B). Unique Interaction tables of proteins significantly enriched in one a sample type compared to a different sample type by SAINT analysis are available in S3 Table. Interestingly, we observed that the extent of self-labeling of Hsp60L-TurboID was roughly 2-fold greater in the soluble fraction relative to the membrane fraction (S3 Table, highlighted in blue). We suggest that this may be due to the relatively ATP-rich environment of the cytoplasm relative to the mitochondrion, thus potentially explaining why we did not see a reduction in labeling of cytosolic proteins as knockdown proceeded (Fig 3C). In examining the SAINT analyses, we identified 122 proteins that were significantly enriched in the membrane fraction relative to both the soluble and no biotin control samples (Fig 4B). We hereafter refer to this set of proteins as a list of “putative mitochondrial” proteins. We retrieved the “MitoProtII” (MPII) [19] and *“Plasmodium* MitoCarta” (PMC) [20] scores for these 122 proteins to ascertain the degree to which the list is enriched in proteins predicted to be mitochondrial. These scores are produced computationally by analyzing N-terminal targeting sequences for the features of mitochondrial targeting peptides (MPII), or by combining multiple factors hypothesized to be common to mitochondrial proteins (PMC). We also obtained scores for a mitochondrial “gold positive” protein set compiled by our group and a non-mitochondrial “gold negative” set of previously published non-mitochondrial proteins [20], and constructed violin plots to compare the distributions of scores (Fig 4C). PMC scoring appeared to provide a more nearly complete separation of the positive and negative sets, although the difference between the average scores of the sets was greater with MPII scoring (S4 Table). Looking at the PMC data for our experimental protein set, we observed that the list of 122 putative mitochondrial proteins had higher scores on average than the “gold negative” list, yet lower than the “gold positive” list, suggesting the list may be incompletely enriched for mitochondrial proteins (Fig 4C). We also conducted literature searches to ascertain which of these 122 had previously been localized to the *P. falciparum* mitochondrion, which had been localized elsewhere, and the gene ontology predicted localization for the remaining proteins. This data is available in S5 Table.

**Fig 4.**
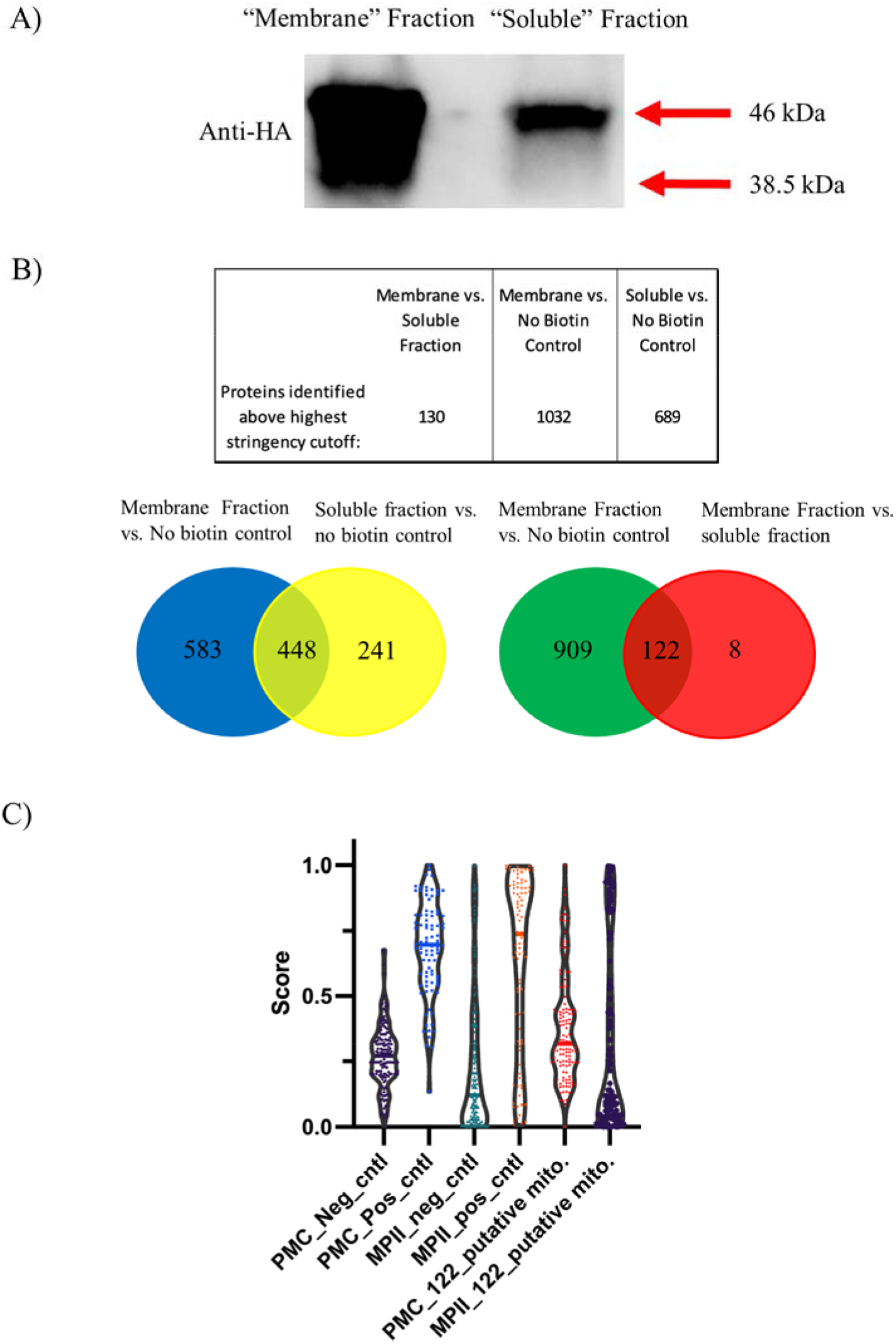
Mass spectrometry results of mitochondrial proximity biotinylation experiments after hypotonic lysis fractionation. A) Western blot analysis of Hsp60L-TurboID-3xHA samples after hypotonic lysis. Note that the soluble fraction only has the band corresponding to full-length Hsp60L-TurboID-3xHA, while the membrane fraction appears to contain a mix of both imported and unimported species. B) LC-MS/MS results analyzed by SAINT analysis (n = 3 biological replicates). Significantly enriched proteins were identified above the highest stringency cutoff. Note the 122 proteins enriched in both membrane fraction vs. no biotin control and membrane fraction vs. soluble fraction. Spectral count data derived from Trans Proteomic Pipeline analysis of all proteins detected in each sample are available in S2 Table. C) Violin plots showing PMC and MitoprotII scores for 3 different sets of proteins. Neg_cntrl and Pos_cntrl are the “gold negative” non-mitochondrial and “gold positive proteins” mitochondrial protein sets described in the text. 122_putative mito. corresponds to the set of proteins generated by analysis of the fractionation experiments in this study. Proteins in each set, PMC and MPII scores, and descriptive statistics for the data set are available in S4 Table.

### Validation of putative mitochondrial proteins identified by proximity biotinylation

To validate our screen, we next wanted to determine if a subset of proteins from our “putative mitochondrial” list generated by our proteomic experiments were indeed localized to the mitochondrion. We selected proteins that had a range of MitoProtII scores, had not been previously localized, and were generally predicted to be essential according to phenotype genome wide disruption studies [21, 22] (Table 1). Using these criteria, we selected five proteins for validation: PF3D7_1237700, PF3D7_0707400, PF3D7_1437700, and PF3D7_0514900, and PF3D7_0105500. Although this last protein was not in the final list of 122, we had identified PF3D7_0105500 early in our pilot experiments with proximity biotinylation and LC-MS/MS as a candidate mitochondrial protein (data not shown).

**Table 1.** Candidate proteins selected for localization study.

| Gene ID<sup>a</sup> | Product description<sup>a</sup> | MitoProtII Score<sup>b</sup> | PMC Score<sup>c</sup> | Mutagenesis index score<sup>d</sup> | IFA result<sup>e</sup> |
| --- | --- | --- | --- | --- | --- |
| PF3D7_1237700 | Conserved protein, unknown function | 0.74 | 0.789 | 0.19 | + |
| PF3D7_0707400 | AAA family ATPase, putative | 0.63 | 0.747 | 0.13 | + |
| PF3D7_1437700 | Succinyl-CoA ligase, putative | 0.72 | 0.585 | 1 | + |
| PF3D7_0105500 | Conserved protein, unknown | 0.99 | 0.674 | 0.161 | + |

|  | function |  |  |  |  |
| --- | --- | --- | --- | --- | --- |
| PF3D7_0514900 | Conserved *Plasmodium* protein, unknown function | .05 | 0.208 | 0.12 | - |

<sup>a</sup>taken from PlasmoDB (plasmodb.org), release 56 [24].
<sup>b</sup>ihg.helmholtz-muenchen.de/ihg/mitoprot.html [19]; scores range from 0 to 1 with higher scores indicating increased likelihood of targeting to the mitochondrion.
<sup>c</sup>Table S1 of van Esveld SL, *et al.* [20]; scores were linearly scaled to range from 0 to 1 for comparison with MitoProtII scores; scores > 0.5416 were considered to indicate likely mitochondrial localization.
<sup>d</sup>Zhang *et al.* [21]; scores range from 0.1 to 1, and the closer the score is to 0.1, the more likely the gene was considered to be essential.
<sup>e</sup>+ indicates predominant co-localization with MitoTracker red in immunofluorescence assays reported in this work.

We generated transgenic parasites in which four of these genes were tagged with 3xHA and expressed from an ectopic location under the control of the moderate mRL2 promoter using the *BxbI attB* x *attP* system [17, 23]. PF3D7_1437700 was endogenously tagged with 3xHA using CRISPR/Cas9 and a donor plasmid containing the TetR-DOZI-Aptamer system [15]. We confirmed that these proteins were expressed *in vivo* at their expected sizes via western blot analysis using an anti-HA antibody (Fig 5). Subcellular localization of these proteins was carried out by indirect immunofluorescence assays. As shown in Fig 6, we observed clear mitochondrial localization for PF3D7_1237700, PF3D7_0707400, PF3D7_1437700, and PF3D7_0105500 (Fig 6). PF3D7_0514900 did not localize with the mitochondrion but displayed punctate like staining throughout the parasite cytoplasm which was consistent with its low MitoProtII score (Table 1).

**Fig 5.**
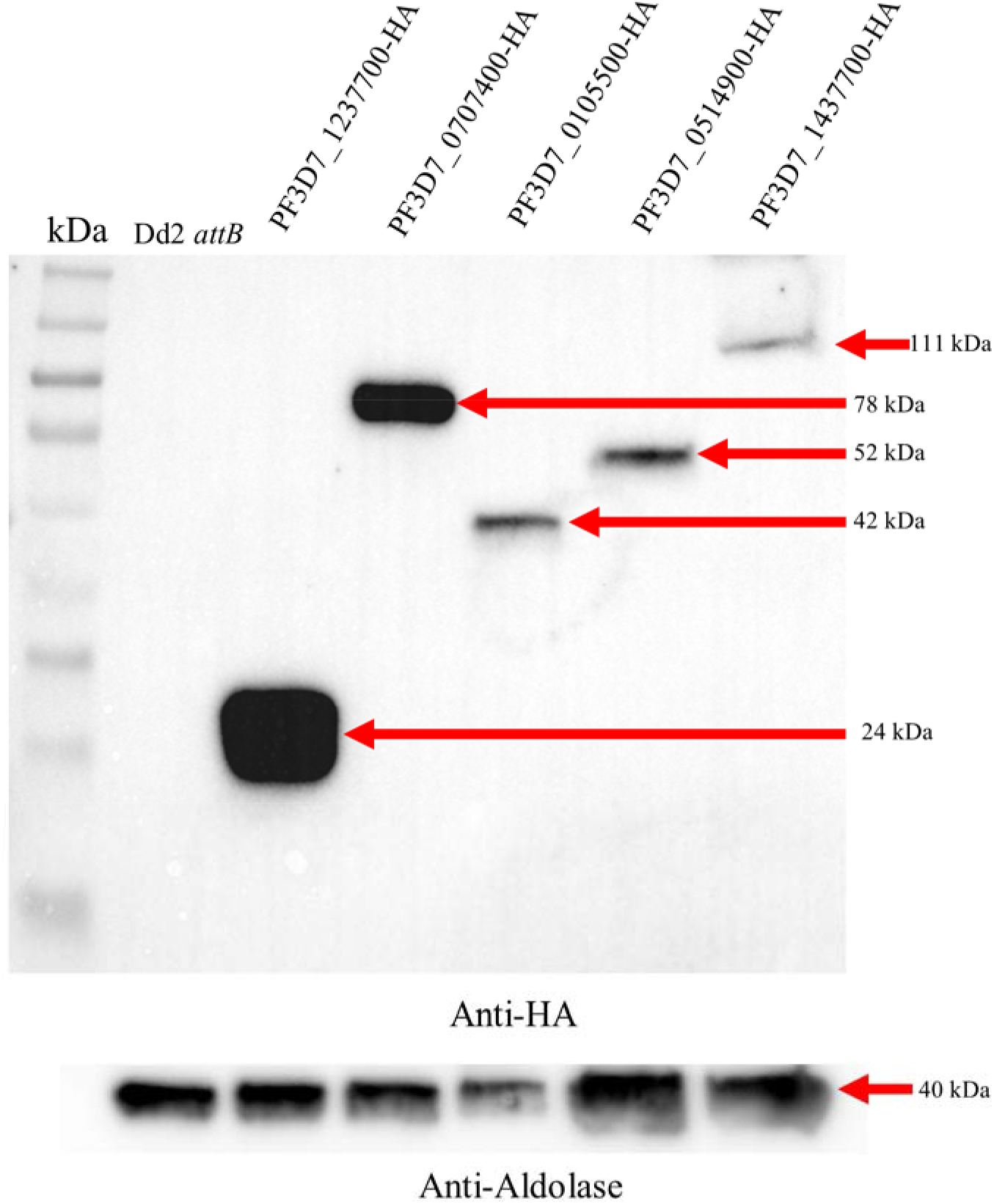
Western blot analysis of parasite lines generated for localization study. The PVDF membrane was probed with anti-HA antibody, stripped, and then probed with an anti-aldolase antibody as a loading control. Expected sizes are as follows: PF3D7_1237700 ~24kDa, PF3D7_0707400 ~78kDa, PF3D7_0105500 ~42kDa, PF3D7_0514900 ~52kDa, PF3D7_1437700 ~111kDa, and aldolase ~41kDa. Colorimetric and chemiluminescent images were merged to include protein molecular weight markers using Image Lab 6.1 (Bio-Rad).

**Fig 6.**
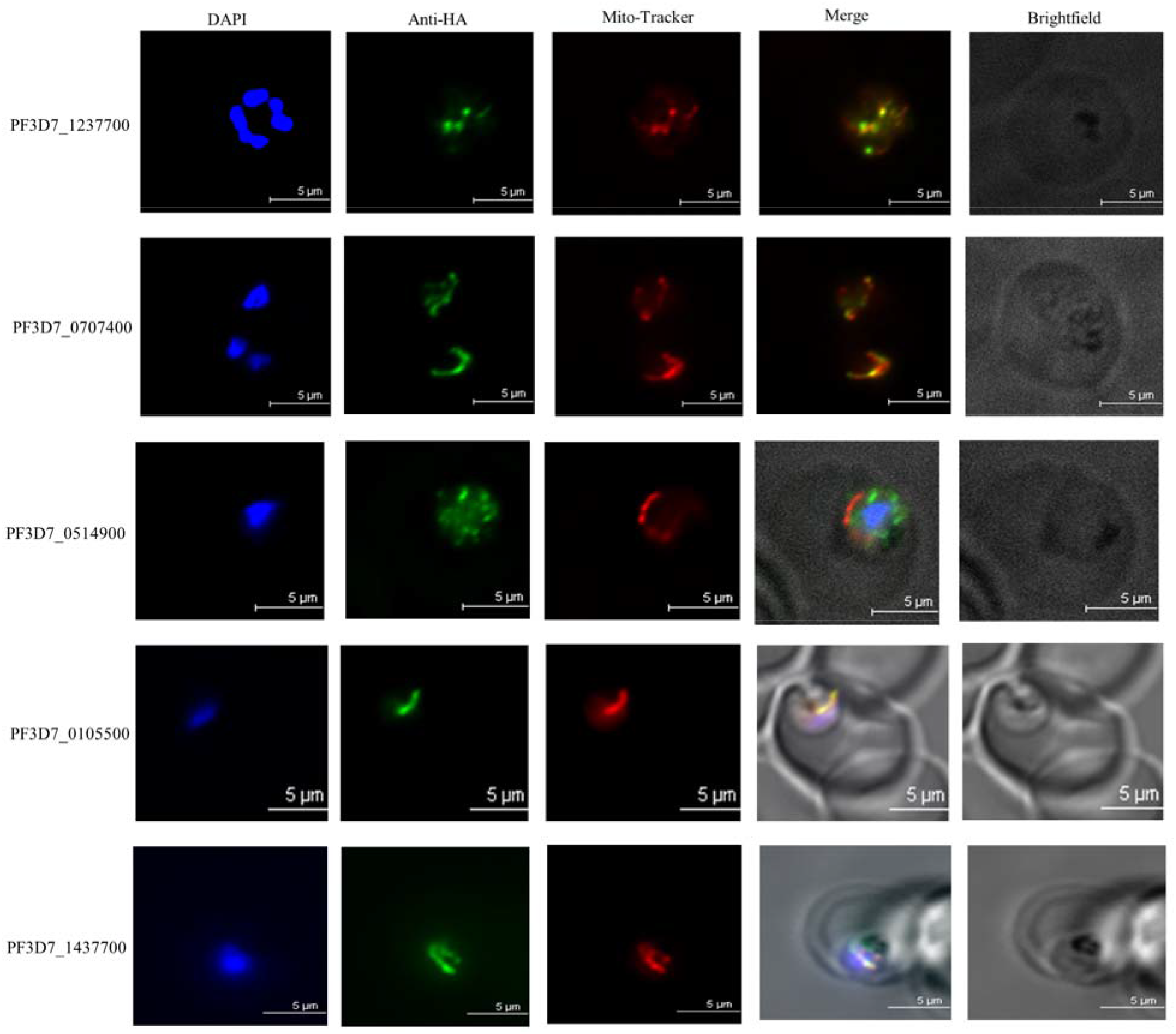
Localization of proteins identified by proximity biotinylation. Immunofluorescence assays were conducted to observe the localization of selected putative mitochondrial proteins. Live parasites were stained with Mito-Tracker (red). Parasites were then fixed and stained with DAPI (blue) and anti-HA antibody (green). Images are representative of ~20 captured images. Scale bars denote 5μm.

## Discussion

Antimalarials that target mitochondrial proteins in asexual blood stage parasites have been therapeutically successful, despite the organelle’s minimal contribution to energy generation at this stage. Due to this success, designing novel therapeutics to target additional *P. falciparum* mitochondrial proteins is an attractive strategy. However, the *P. falciparum* mitochondrial proteome has yet to be fully defined. During analysis of the initial *P. falciparum* genome annotation, we predicted at least 148 nuclear-encoded proteins were targeted to the mitochondrion based upon homology [25]. Bender *et al*. analyzed the N-terminal sequences of putative *P. falciparum* mitochondrial proteins and developed a neural network program to predict mitochondrial targeting of *P. falciparum* proteins, which predicted between 381 (7%) to 1,177 proteins (22%) to be targeted to the mitochondrion, depending on the stringency of the network [26]. A more recent study analyzed eight proteomic data sets relevant to mitochondrial protein expression and targeting, and employed Bayesian statistics to generate a list of 445 putative mitochondrial proteins in *P. falciparum* [20]. While useful, these predicted sets of mitochondrial proteins were not experimentally derived and are observed to contain a certain number of misassignments that varies with the particular analysis and its parameters. On the other hand, proximity biotinylation experiments were used successfully in another apicomplexan parasite, *Toxoplasma gondii*, identifying 421 putative mitochondrial proteins that included a large number of known mitochondrial proteins in *T. gondii* [27].

In this study, we employed a proximity biotinylation strategy with TurboID fused to the leader sequence to that was also used in a study conducted in *T. gondii* (*Pf*Hsp60L vs. *Tg*Hsp60L) to generate an experimental set of putative *P. falciparum* mitochondrial proteins. Despite trying multiple different experimental protocols, we were able to generate a statistically significant set of 122 putative mitochondrial proteins, prompting us to contemplate possible reasons for such a low number of proteins to be identified. One possibility is that Hsp60L is sub-optimal for mitochondrial targeting in *P. falciparum*. Although *Pf*Hsp60L has been previously used for targeting proteins to the mitochondrion [16], a more extensive investigation has shown an accumulation of native *Pf*Hsp60 in the cytosol in asexual blood stage parasites as well as being imported to the mitochondrion [28]. It is therefore possible that PfHsp60L-TurboID also persists in the cytosol, despite the apparent lack of significant cytosolic fluorescence in the immunofluorescence assay (Fig 2B). Our observation of continued presence both precursor and processed forms of Hsp60L-TurboID even after its expression was suppressed (Fig 3B) is consistent with this notion of Hsp60 being equally redistributed between the cytosol and mitochondrion. These results are very different from the study utilizing *Tg*Hsp60L-BirA* in proximity biotinylation experiments of the mitochondrial matrix, in which the lower molecular weight processed protein was dramatically more intense than the full-length, unprocessed protein [27]. This suggests a more efficient processing and/or import into the mitochondrion of *T. gondii* compared to *P. falciparum*. Future experiments will be required to determine if this inefficient mitochondrial import is a general feature of *P. falciparum* relative to other organisms, or if this phenomenon is limited to the *Pf*Hsp60 leader sequence. Though not mutually exclusive, there is also the possibility that the asexual blood stage *P. falciparum* mitochondrion does not have sufficient ATP required to furnish robust biotinylation of proteins, a reaction that requires ATP. Although subunits of ATP synthase are essential for the survival of asexual stage parasites, the mitochondrion is not a major source of ATP, unlike in *T. gondii* [21, 23]. This may explain why we observed self-labeling of Hsp60L-TurboID to be roughly 2-fold greater in the soluble fraction compared to the membrane fraction (S3 Table, highlighted in blue), and more generally the challenges of applying proximity biotinylation techniques to the mitochondrion of asexual blood stages of *P. falciparum*. Development of tools to rigorously monitor organellar ATP levels in *P. falciparum* and other Apicomplexans will be necessary to test this possibility.

Even from our “putative mitochondrial” candidate list, we were able to successfully localize four proteins of unknown function by immunofluorescence experiments (Fig 6) that have not, to our knowledge, been previously localized. Non-mitochondrial localization of PF3D7_0514900 is not altogether surprising, as this protein had a very low mitochondrial targeting probability (Table 1) compared to the others selected for validation. The mitochondrial localization of these proteins in some cases was unexpected. Previous studies have reported PF3D7_1237700 as being likely nuclear [29] or predicted it to be localized to the surface of sporozoites [30]. We show clear mitochondrial localization of this protein by indirect immunofluorescence experiments in blood stage parasites (Fig 5). This protein has also been shown to be likely essential by a published *piggyBac* saturation mutagenesis screen [21], but has no annotated function. Further studies are needed to elucidate the physiological contributions of PF3D7_1237700 and to assess its potential as a therapeutic target. The *piggyBac* saturation mutagenesis screen also demonstrated PF3D7_0707400 as being likely essential (Table 1). This gene is annotated as a putative AAA+ ATPase, and appears to bear orthology to the well-studied mammalian mitochondrial protein ATAD3A. Recently, this protein has been assigned multiple critical functions in mammals. These include: cristae formation [31, 32], mitochondrial nucleoid stabilization [33], mtDNA replication [32], and mitochondrial translation [33]. Mutations and splice variants in human ATAD3A have also been implicated in numerous genetic disorders [34, 35], as well as in cancers [36] and neurological syndromes [37]. Presence of a potential ATAD3A orthologue in a highly divergent organism like *P. falciparum* demands further investigations of its role in mitochondrial physiology of the parasite. In contrast, the gene product of PF3D7_0105500 localized to the mitochondrion appears restricted only to *Myzozoans*, a sub-group of phyla within the *Alveolata* clade. The fact that this gene product is so divergent from host proteins, combined with its likely essentiality, makes it an attractive candidate for an antimalarial drug target. Detailed phenotypic characterization of this mitochondrial protein is also likely to reveal evolutionary adaptations within this group of deeply branching eukaryotes.

## Methods

### Immunofluorescence assays

Fifty microliters of packed volume parasite cultures were pre-labeled with 60nM MitoTracker Red CMXRos (Life Technologies by ThermoFisher Scientific) for 30 min at 37°C. Samples were then washed 3 times with PBS. Cells were then fixed with 4% v/v paraformaldehyde and 0.0075% v/v glutaraldehyde for 1 h at 37°C with agitation. Cells were washed three times with PBS and permeabilized with 0.5% v/v Triton X-100 in PBS for 10 min on a rotor. Cells were next incubated with NaBH4 (0.1 mg/ml) for 5 min at room temperature on a rotor and blocked with 5% w/v BSA overnight at 4°C. Parasites were then incubated with the HA probe (sc-7392, Santa Cruz Biotechnology) at a 1:250 dilution overnight with agitation at 4°C. After three washes with PBS, the secondary antibody, Alexa Fluor 488 conjugated anti-mouse IgG (Life Technologies by ThermoFisher Scientific), was added at a 1:250 dilution overnight at 4°C. Parasites were then washed 3 times with PBS, resuspended in anti-fade buffer (S2828, ThermoFisher Scientific), and visualized using a Nikon Ti microscope.

### Plasmid construction

To generate an aTc-regulated Hsp60L-TurboID-3xHA plasmid for transfection, the pLN-RL2-*hdhfr*-Hsp60L-mNeonGreen-3xHA plasmid (a gift from Dr. Hangjun Ke, Drexel University) was obtained as a backbone. PCR amplification of TurboID from TurboID-his bacteria was performed to add a 5’ NheI site and a 3’ AflII site. This fragment was then inserted in place of mNeonGreen, yielding pLN-RL2-Hsp60L-Turbo-ID-3xHA. The Hsp60L-Turbo-ID fusion protein was amplified by PCR, adding a 5’ EcoRV site and a 3’ BsteII site. This product was then ligated into digested and gel purified pMG75-*attP*, yielding pMG75-*attP*-Hsp60L-TurboID-3xHA. For localization studies, the coding region of PF3D7_0105500, PF3D7_0707400, and PF3D7_0514900 was cloned into the pLNmRL2 vector bearing a 3xHA tag by PCR [38]. The coding region of PF3D7_1237700 was synthesized by Genewiz (Azenta Life Sciences) and cloned into the pLNmRL2 vector.

### Parasite lines, culture, and transfection

Dd2 *attB P. falciparum* WT parasites were used in this study. Asexual *P. falciparum* parasites were maintained as previously described [39]. Briefly, parasites were cultured with RPMI 1640 medium supplemented with 0.5% w/v Albumax II (Gibco by ThermoFisher Scientific), sodium bicarbonate (2.1g/liter, Corning by ThermoFisher Scientific), HEPES (15 mM, MilliporeSigma), hypoxanthine (10mg/liter, Fisher Scientific), and gentamycin (50mg/liter, VWR). Prior to transfection, pLN vectors were co-precipitated with the integrase plasmid using 3M Potassium/5M Acetate pH 7 and ethanol. Transfections were done by electroporation via a Bio-Rad gene pulser (0.31 kV, 960 μFD). Electroporation was performed on cultures of 5-6% rings mixed with 40μg of circular pLN plasmid and 40μg circular integrase plasmid dissolved in cytomix (120 mM KCl; 0.15 mM CaCl_2_, 10 mM K_2_HPO_4_/KH_2_PO_4_, pH 7.6; 25 mM Hepes, pH 7.6; 2 mM EGTA, pH 7.6; 5 mM MgCl_2_). Transfections were carried out as previously described [40]. After electroporation, parasites were cultured under standard conditions for 48 hours before the addition of blasticidin (2.5μg/ml, InvivoGen). Unless otherwise specified cultures were maintained *in vitro* below a parasitemia of 5%.

### Western blotting

Parasite cultures were treated with 0.05% w/v saponin in PBS supplemented with a 1X protease inhibitor mixture (P8215, MilliporeSigma). The remaining pellet was resuspended in ~10 volumes of 2% w/v SDS, 62mM Tris-HCl (pH 6.8) and mixed by vigorous pipetting. The samples were then mixed by rotation at room temperature. This lysate was then spun down at max speed for 5 min in a clinical centrifuge, and the resulting supernatant was used for SDS-PAGE. Membranes blotted with streptavidin-HRP (Invitrogen) were blocked with 5% w/v dry milk, while membranes blotted with anti-HA antibodies were blocked with 5% w/v BSA. Blots were then incubated with a mouse monoclonal anti-HA antibody (sc-7392, Santa Cruz Biotechnology) at 1:12,500 and a horseradish peroxidase-conjugated secondary goat anti-mouse antibody (A16078, ThermoFisher Scientific) at 1:12,500. For detection of biotinylated proteins, membranes were incubated with 1:20,000 streptavidin-HRP for 2 hours at room temperature. The blots later probed with a rabbit anti-*Pf*Exp2 primary antibody (a gift from Dr. James Burns, Drexel University) at 1:10,000 followed by a horseradish peroxidase-conjugated secondary goat anti-rabbit antibody (31460, ThermoFisher Scientific) at 1:20,000. All other steps followed the standard protocol.

### Proximity biotinylation fractionation assay

Prior to the experiment, cultures were grown in biotin-free medium (USA Biologicals) and synchronized with two volumes of 0.3M alanine buffered with 10mM HEPES (pH 7.5). For each of the three “no biotin control” samples, 2.5ml of packed ~8% trophozoites were grown in T-175 flasks. For each of the three samples that were subsequently fractionated into membrane and soluble fractions, 5ml of packed ~8% trophozoites in two T-175 flasks were used. On the day of the experiment, cultures were treated with 150μM of biotin dissolved in 95% v/v ethanol for 2 hours, except for the three “no biotin control” flasks. All parasites were then released by treatment with 0.05% w/v saponin in PBS. No biotin control samples were then solubilized in RIPA buffer containing protease inhibitors for 1 hour at room temperature on a rotor, and then incubated on ice for 20 min. The remaining samples were fractionated by hypotonic lysis and low-speed centrifugation as previously described [28]. Five hundred microliters of streptavidin-conjugated magnetic beads (Invitrogen Dynabeads MyOne Streptavidin C1) were washed with PBS and added directly to the three “soluble fraction” samples and rotated overnight at 4°C. The three “membrane fraction” samples were then solubilized with RIPA buffer containing protease inhibitors and rotated at room temperature before incubation on ice for 20 min. The six RIPA-solubilized no biotin and membrane fraction samples were then subjected to two rounds of sonication (30 seconds with 30% on, 70% off duty cycle) on a Fisher Scientific Sonic Dismembrator (model 500) using a 1/8” microtip. Remaining insoluble material was removed by centrifugation at 14,000xg. Five hundred microliters of streptavidin-conjugated magnetic beads were added to these six samples, and they were rotated overnight at 4°C. All beads were then washed with wash buffers 1-3 [41, 42] for 10 minutes each, washed ten times with 50mM ammonium bicarbonate, and frozen at −80°C for on-bead digestion and LC-MS/MS analysis.

### On-bead digestion

Trypsin (0.2ug in 20 ul 50mM NH_3_HCO_4_) was added to washed and ready to digest beads and incubated at 37C for 4 hours. Another 0.2 ug of trypsin was added and incubated at 37C overnight. The solution was separated from the magnetic beads and pH was adjusted to 3 with 10% formic acid. The sample was desalted with stage tip before LC-MS/MS.

### Liquid chromatography-tandem mass spectrometry (LC-MS/MS)

Samples were analyzed by LC-MS using Nano LC-MS/MS (Dionex Ultimate 3000 RLSCnano System) interfaced with Orbitrap Eclipse Tribrid (Thermofisher). Samples were loaded on to a fused silica trap column Acclaim PepMap 100, 75umx2cm (ThermoFisher). After washing for 5 min at 5 μl/min with 0.1% v/v TFA, the trap column was brought in-line with an analytical column (Nanoease MZ peptide BEH C18, 130A, 1.7um, 75umx250mm, Waters) for LC-MS/MS. Peptides were fractionated at 300 nL/min using a segmented linear gradient 4-15% B in 30min (where A: 0.2% v/v formic acid, and B: 0.16% v/v formic acid, 80% v/v acetonitrile), 15-25%B in 40min, 25-50%B in 44min, and 50-90%B in 11min. Solution B then returns at 4% for 5 minutes for the next run.

The scan sequence began with an MS1 spectrum (Orbitrap analysis, resolution 120,000, scan range from M/Z 350–1600, automatic gain control (AGC) target 1E6, maximum injection time 100□ms). The top S (3 sec) and dynamic exclusion of 60sec were used for selection of Parent ions for MSMS. Parent masses were isolated in the quadrupole with an isolation window of 1.4 m/z, automatic gain control (AGC) target 1E5, and fragmented with higher-energy collisional dissociation with a normalized collision energy of 30%. The fragments were scanned in Orbitrap with resolution of 30,000. The MS-MS scan range was determined by charge state of the parent ion, but lower limit was set at 100 amu.

### Database Search

LC-MS/MS data was processed using the Trans-Proteomic Pipeline (TPP) [43], as previously described [44]. Raw mass spectra were converted to .mzML format using MSConvert [45]. Spectra were searched using X!Tandem and Comet against reference proteomes for *P. falciparum* Pf3D7 (PlasmoDB, v53), human (Uniprot), common contaminants (Contaminant Repository for Affinity Purification Mass Spectrometry Data) [46], and randomized decoys generated through TPP. X!Tandem and Comet searches were combined in iProphet and proteins were identified with ProteinProphet and proteins were identified with ProteinProphet. Proteins identified in this search below a 1% false positive error rate are reported. Biological replicates were combine and compared across samples using SAINT version 2.5.0 [18]. Proteins with a SAINT score <0.1 were considered enriched, as previously used for high stringency significance [47].

## Acknowledgements

We thank all members of the Vaidya lab for lively discussion and input. This work was supported by the National Institutes of Health grant 5R01AI028398-31 awarded to A.B.V., by an R01 from the National Institutes of Allergy and Infectious Diseases (R01AI123341 to SEL), and by funding in support of KTR by the Pennsylvania State University (COVID Relief Funds).

## Supporting Information

**S1 Table. Spectral counts for each protein identified when Hsp60L-TurboID was knocked down for different amounts of time.** This table contains spectral count data derived from Trans Proteomic Pipeline analysis of the experiment described in Fig 3.

**S2 Table. Spectral counts for each protein identified after fractionation by hypotonic lysis.** This table contains the spectral count data derived from Trans Proteomic Pipeline analysis of the experiment described in Fig 4.

**S3 Table. Unique Interaction tables of SAINT analysis outputs for comparisons between the membrane fraction, soluble fraction, and no biotin control.** The different tabs in the table indicate proteins enriched in a sample compared to another sample as determined by SAINT analysis of Trans Proteomic Pipeline output.

**S4 Table. MPII and PMC scores for three protein sets: 122_putative mito., gold negative, and confirmed positive.** This table Includes MitoProtII and *Plasmodium* MitoCarta scores in the three protein sets, which are predicted likelihoods of mitochondrial targeting. Descriptive statistics for these scores are also included.

**S5 Table. Results of literature searches for published localization data on the 122_putative mito. protein set.** Proteins localized to the mitochondrion are highlighted in green. Proteins localized elsewhere are highlighted in red. Highlighted in blue are the proteins localized in this paper.

